# Desiccation-induced changes in recombination rate and crossover interference in *Drosophila melanogaster*: evidence for fitness-dependent plasticity

**DOI:** 10.1101/382259

**Authors:** Dau Dayal Aggarwal, Sviatoslav R. Rybnikov, Irit Cohen, Zeev Frenkel, Eugenia Rashkovetsky, Pawel Michalak, Abraham B. Korol

## Abstract

Meiotic recombination is evolutionarily ambiguous, as being associated with both benefits and costs to its bearers, with the resultant dependent on a variety of conditions. While existing theoretical models explain the emergence and maintenance of recombination, some of its essential features remain underexplored. Here we focus on one such feature, recombination plasticity, and test whether recombination response to stress is fitness-dependent. We compare desiccation stress effects on recombination rate and crossover interference in chromosome 3 between desiccation-sensitive and desiccation-tolerant *Drosophila* lines. We show that relative to desiccation-tolerant genotypes, desiccation-sensitive genotypes exhibit a significant segment-specific increase in single- and double-crossover frequencies across the pericentromeric region of chromosome 3. Significant changes (relaxation) in crossover interference were found for the interval pairs flanking the centromere and extending to the left arm of the chromosome. These results indicate that desiccation is a recombinogenic factor and that desiccation-induced changes in both recombination rate and crossover interference are fitness-dependent, with a tendency of less fitted individuals to produce more variable progeny. Such a dependence may play an important role in the regulation of genetic variation in populations experiencing environmental challenges.

## INTRODUCTION

Since the seminal experiments by Plough (Plough 1917, 1921), meiotic recombination has been known to possess plasticity with respect to different ecological factors. Typically, its rates are lower in an optimal environment, and rise when conditions become more stressful. To date, the plasticity of recombination rate has been reported for different species and various ecological factors (for recent reviews see (Bomblies et al. 2015; Modliszewski and Copenhaver 2017; Stapley et al. 2017)). Notably, this phenomenon was observed in fruit flies with respect to heat (Plough 1917, 1921; Stern 1926; Graubard 1932; Politzer 1940; Hayman and Parsons 1960; Chandley 1968; Grell 1978; Korol et al. 1994; Zhong and Priest 2011; Jackson et al. 2015), cold (Plough 1917; Graubard 1932; Politzer 1940; Zhong and Priest 2011), starvation (Neel 1941), specific chemicals (Kilias et al. 1979), mating stress (Zhong and Priest 2011), and parasite infection (Singh et al. 2015; Singh 2019), although higher production of recombinant offspring may have resulted from a transmission distortion rather than elevated recombination rates in the last case (Singh et al. 2015). In their experiments with tomato plants, Korol *et al.* demonstrated that chiasma numbers grew in cold-resistant genotypes under the high-temperature cultivation regime, while in heat-resistant ones – under the low-temperature regime (Zhuchenko et al. 1986). Based on these results, the authors suggested that the plasticity of recombination rate is modulated by genotype fitness, which can be manifested as a negative recombination-fitness association. However, such *inter-genotype* studies remain extremely limited. Particularly, in fruit flies plasticity of recombination rate was shown to be fitness-dependent only with respect to heat stress (Zhuchenko and Korol 1985; Korol et al. 1994; Zhong and Priest 2011) and specific chemicals associated with oxidative stress (Kilias et al. 1979; Hunter et al. 2016).

One of the essential recombination features is crossover interference, when the frequency of double crossovers may appear either lower (positive interference) or higher (negative interference) than that the expected under the assumption of crossover independence. The phenomenon and its evolutionary significance attract increasing interest (Segura et al. 2013; Bomblies et al. 2016), even though variation in interference across environments and genotypes remains underexplored. To date, interference plasticity in fruit flies was reported only with respect to heat, associated with a consistent increase in double-crossover frequency under stress (Plough 1921; Graubard 1934; Hayman and Parsons 1960; Grell 1978). However, the question of whether plasticity of interference is fitness-dependent has never been studied, to the best of our knowledge.

In this study, we aimed to test if desiccation-induced changes in recombination rate and interference depend on flies’ desiccation tolerance. We used *D. melanogaster* lines with differential desiccation tolerance that have recently been established as a part of our long-term experiment aimed at testing whether directional selection for desiccation tolerance may cause indirect selection for recombination (Aggarwal et al. 2015). In that study, we have found, in accordance with theoretical predictions (Charlesworth 1993), that the selected lines evolved, along with the increased desiccation tolerance, a segment-specific increase in recombination rate compared to the non-selected, desiccation-sensitive ones. We also observed a segment-specific relaxation of positive crossover interference (and even appearance of negative interference) in the selected lines, and these changes were not necessarily coupled with changes in recombination rate. In the current study, we assumed and explicitly confirmed that the difference between selected and non-selected lines in desiccation tolerance holds also for their F_1_ hybrids with a standard multiple-marker strain. This allowed us to use F_1_ heterozygous females for testing the hypothesis that desiccation-sensitive genotypes display higher plasticity of recombination rate and interference with respect to desiccation stress, compared to desiccation-tolerant ones, consistent with the concept of condition-dependent recombination.

## MATERIAL AND METHODS

### Flies and experimental arrangement

We used two types of parental lines: (a) desiccation-sensitive (*S*), originating from flies collected in 2009 in Madhya Pradesh, Jabalpur, India; (b) desiccation-tolerant (*T*), originating from the same Jabalpur stock, though underwent long-term (48 generations) laboratory selection for desiccation tolerance (Aggarwal et al. 2015). Virgin females from the parental lines (three *S*- and three *T*-lines, each with three technical replicates; total 18 replicates) were mated with males from the marker stock *ru-cu-ca*, homozygous for six recessive mutations in chromosome 3: *ru, h, th, cu, sr*, and *e*. The mated females were then transferred to fresh food bottles for six hours to lay F_1_ eggs. The third-instar F_1_ larvae were either subject to desiccation treatment or maintained as control (80 larvae per replicate). In *Drosophila*, changes in recombination rate can be induced by exogenous factors during a rather long period – from interphase till the middle-end of pachytene (Chandley 1968; Grell 1978). Three-day-old F_1_ females obtained from both treated and non-treated F_1_ larvae (20 females sampled from each replicate) were back-crossed with males from the same marker line (Figure 1). Recombination and interference were analyzed based on marker segregation in the obtained progeny (750 flies per each of the three S- and three T-lines for the treatment and same for the control variants). Recombination was measured as crossover frequency, while interference – as the coefficient of coincidence, which is the ratio of the observed number of double crossovers to their number expected under the assumption of independent recombination in adjacent intervals.

**Figure 1:**
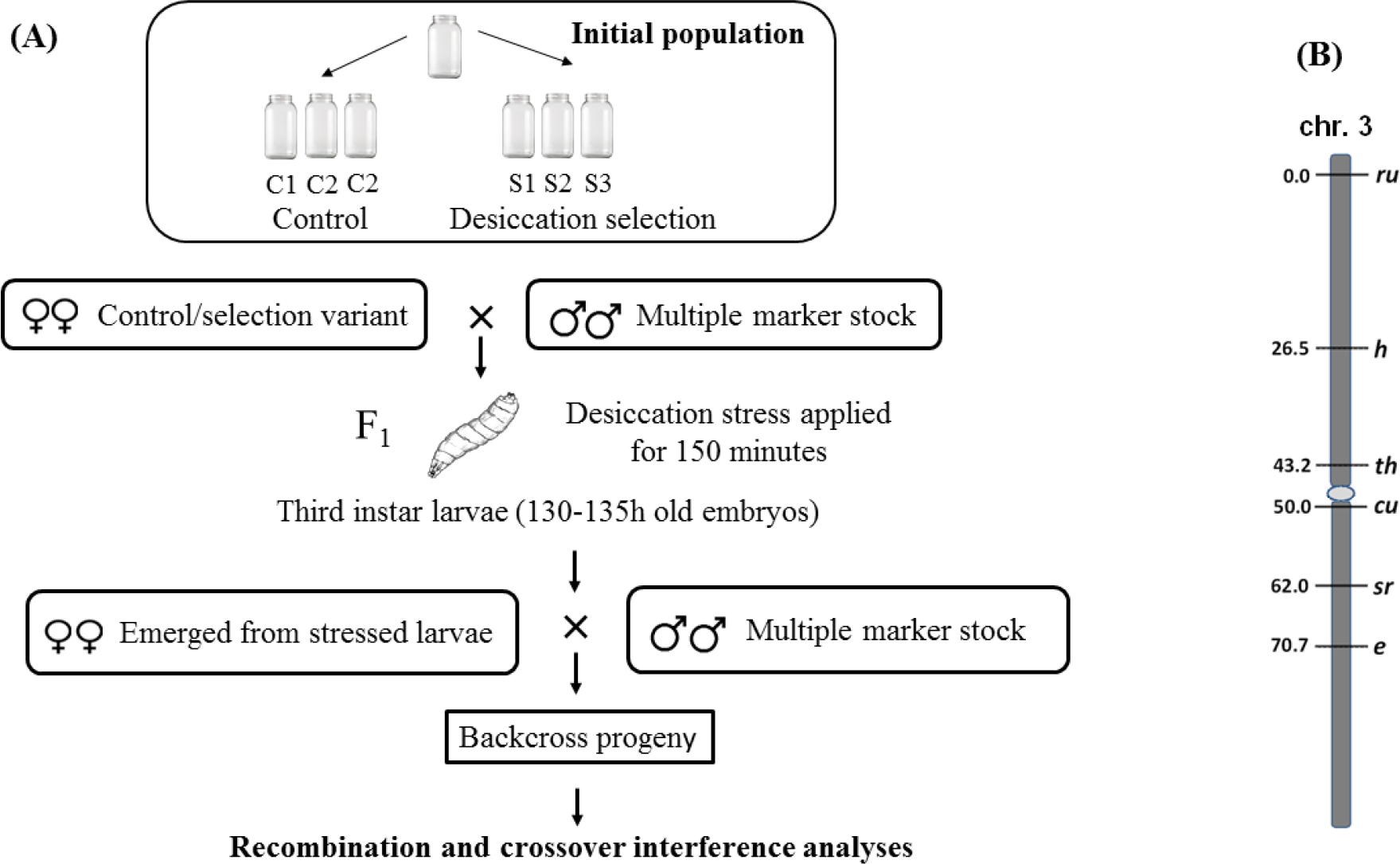
Experimental arrangement (A) and the location of the markers in chromosome 3 (B)

### Treatment and survival analysis

In each of the 18 replicates involved in the experiment, 80 F_1_ larvae were exposed to desiccation treatment and 80 F_1_ larvae were maintained as control. For desiccation treatment, the larvae were divided into 16 groups, each of five larvae (in line with the method by Markow *et al.*, 2007). The blue indicating silica gel (2g) was placed into 16 dry plastic vials (20×90 mm). Groups of 5 larvae, gently isolated with a brush, rinsed, and air-dried, were placed into other 16 empty plastic vials, with their open ends covered with a muslin cloth to enable free air flow. Then, the larvae-containing vials were carefully placed above the silica gel-containing ones, and 16 obtained setups were made airtight by sealing the gaps between the two halves with multi-layers Parafilm. The larvae were treated during 150 min and then transferred into fresh food-containing vials. The F_1_ females hatched from the desiccation-treated larvae (20 females per replicate) were further used in backcrosses. The control flies were reared under the same conditions, except the desiccation treatment. A similar experimental scheme was used in the survival analysis of F_1_ larvae. Groups of 10 larvae were treated by desiccation until death. Groups of the control larvae were left untreated. The number of immobile larvae was scored every 30 min.

### Statistical analysis

For each pair of intervals, maximum likelihood (ML) analysis was performed to estimate the vector of parameters *θ* = {recombination frequencies for the two intervals *r*_1_ and *r*_2_ and the coefficient of coincidence *c*}, for non-treated (control) and treated heterozygous females (*θ*_*n–tr*_ and *θ*_*tr*_, respectively) and to test for significance of the effects of treatment on recombination parameters. For each of the two groups of lines, sensitive (*S*-lines) and tolerant (*T*-lines), the log-likelihood function had the following form:

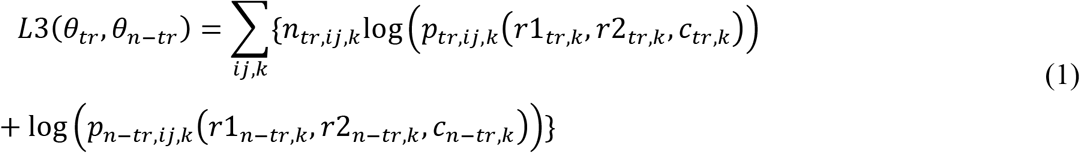

Here *i, j* ∊ {0,1} define whether the recombination event occurred in the first and second interval, respectively (0 - no recombination, 1 - recombination); *n*_*ij*_ represents the observed number of individuals of the genotype class *ij* in the backcross progeny of the given treated or non-treated line *k* (*k*=1,2,3); and *p*_*ij*_ is its expected frequency represented as a function of the unknown parameters:

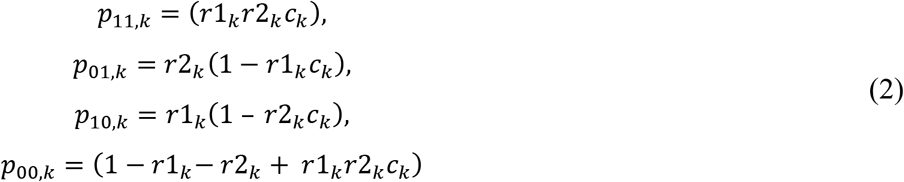

The designation *L*3 for the log-likelihood function was employed here to indicate that the analysis included simultaneously all three recombination parameters per each variant: *r*1, *r*2, and *c*. Yet, for testing the hypotheses about the effect of treatment on recombination rate, we employed log-likelihood function for single-interval analysis:

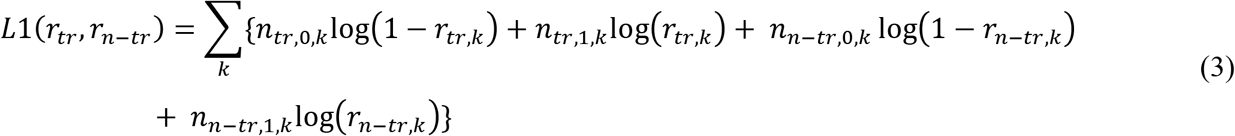

where *n*_*\*,0,k*_ and *n*_*\*,1,k*_ represent the number of observed non-crossover and crossover genotypes in the analyzed progeny of the considered treated or non-treated females. Moreover, this *L*1 function was also employed, in combination with *L*3, for testing the hypotheses about the effect of treatment on interference. ML estimates of the vectors *θ*_*k*_ = (*r*1_*k*_, *r*2_*k*_, *c*_*k*_) for *k*=1,2,3 were obtained by numerical optimization of the log-likelihood function *L*(*θ*_*k*_), using the gradient descent procedure in which all three parameters *r*1_*k*_, *r*2_*k*_ and *c*_*k*_ are evaluated simultaneously in every iteration:

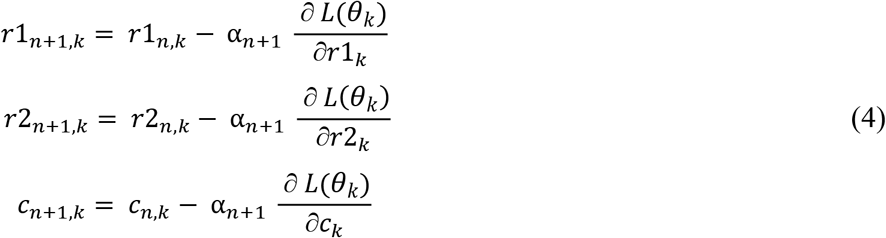

The foregoing joint analysis of the progeny of non-treated and treated F1 females of either tolerant or sensitive lines can be easily extended to include all variants in one log-likelihood ratio test. Thus, to compare the effect of desiccation treatment on recombination rate in desiccation-tolerant (*T*) and desiccation-sensitive (*S*) lines, we examined for each of the five intervals the following hypotheses:

- H_0_ – no difference in recombination rate between the treated and non-treated F_1_ females in both the *S*- and *T*-lines 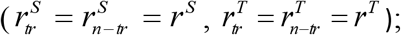
- H_1_ – no difference in recombination rate between the treated and non-treated F_1_ females in the *T*-lines 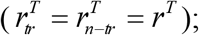
- H_2_ – no difference in recombination rate between the treated and non-treated F_1_ females in the *S*-lines 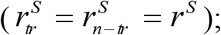
- H_3_ – no restriction on the parameters of the two groups of lines.

Here subscripts *tr* and *n-tr* denote treated and non-treated variants, respectively. To compare H_1_, H_2_ and H_3_ versus H_0_, the log-likelihood ratio test was employed.

For H_1_ versus H_0_,

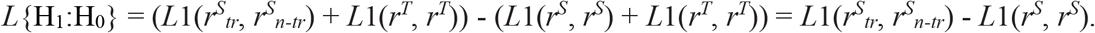

For H_2_ versus H_0_,

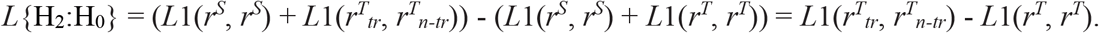

For H_3_ versus H_0_,

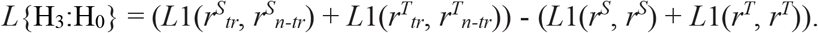

The doubled log-likelihood ratio is distributed asymptotically as chi-square with *df*=3 for the first two tests and *df*=6 for the third test.

Similarly, to compare the effect of desiccation treatment on crossover interference in the *S* and *T* lines, we examined the following hypotheses:

- H_0_ – no difference in crossover interference between the treated and non-treated F_1_ females in both the *S*- and *T*-lines 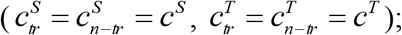
- H_1_ – no difference in crossover interference between the treated and non-treated F_1_ females in the *T*-lines 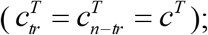
- H_2_ – no difference in crossover interference between the treated and non-treated F_1_ females in the *S*-lines 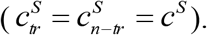
- H_3_ – no restriction on the parameters of the two groups of lines.

For H_1_ versus H_0_,

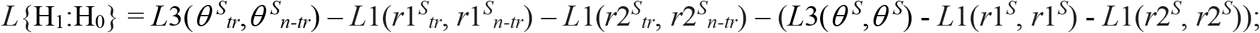

for H_2_ versus H_0_,

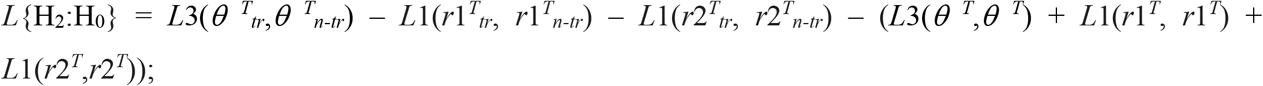

for H_3_ versus H_0_,

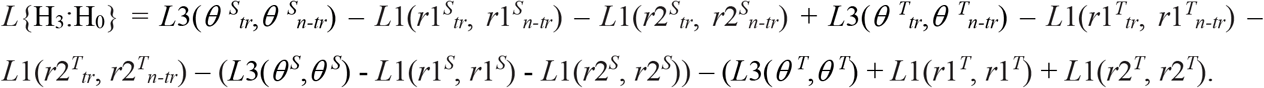

As before, the tests statistics are distributed asymptotically as chi-square with *df*=3 in the first two tests and *df*=6 in the third test.

Gene Ontology enrichment tests for genes found in the *h-th* and *cu-sr* genomic intervals were conducted by contrasting the gene lists from the intervals with all *D. melanogaster* genes using *DAVID* with the Benjamini-Hochberg procedure (Huang et al. 2009).

## RESULTS

Adult flies from the *T*-lines were previously shown to have significantly higher desiccation tolerance compared to those from the parental *S*-lines (Aggarwal et al. 2015; Kang et al. 2016). We assumed that this difference holds also at the larval level, and passes through generations; however, these assumptions had to be tested explicitly. We found that larvae from the parental *T*-lines were indeed significantly more tolerant compared to those of the parental *S*-lines: 1.56-fold increase for LT_100_ (*F*_1, 282_=930.85, *P*<0.0001), and 1.57-fold increase for LT_50_ (*F*_1, 282_=709.86, *P*<0.0001), where LT_100_ and LT_50_ are times to 100% and 50% levels of mortality under dry air, respectively (Figure 2, A). The F_1_ larvae obtained from the parental lines in crosses with the standard multiple-marker strain *ru cu ca* demonstrated the same pattern: 1.42-fold increase for LT_100_ (*F*_1, 282_=729.78, *P*<0.0001), and 1.34-fold for LT_50_ (*F*_1, 282_=582.04, *P*<0.0001) (Figure 2, B). These results indicate that using the terms *S* and *T* is valid not only for the parental lines but also for their F_1_ hybrids with the marker strain.

**Figure 2:**
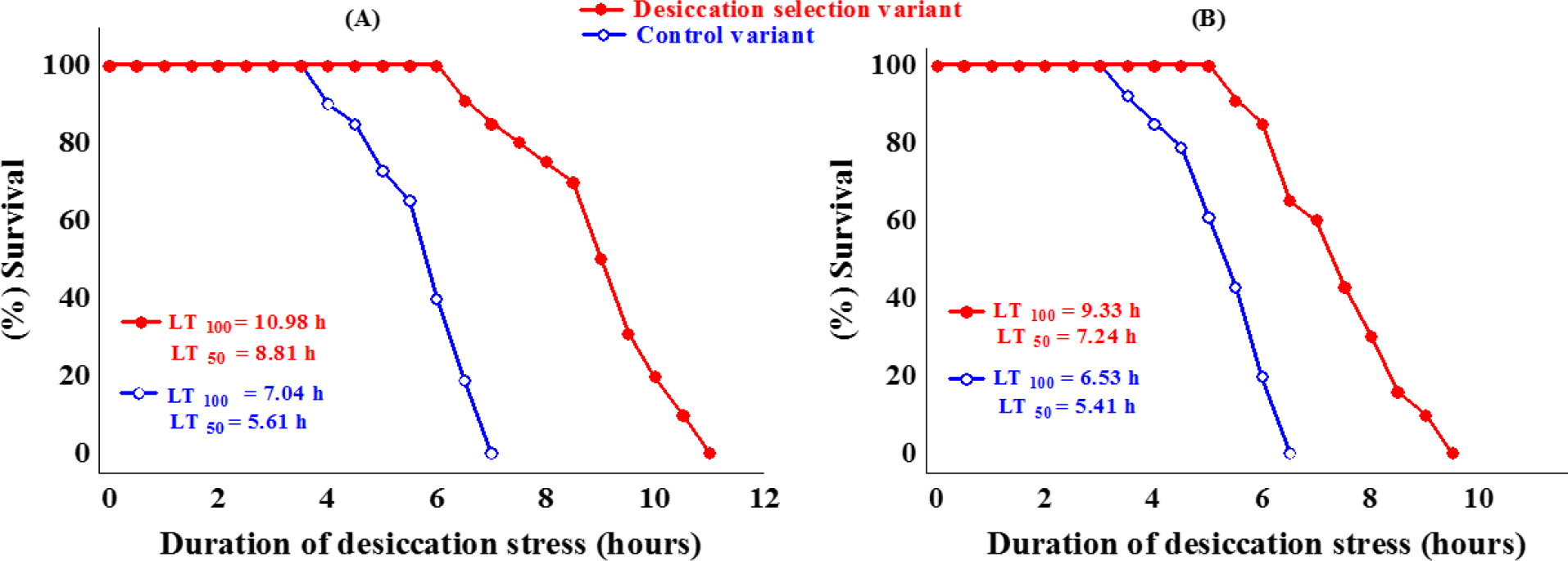
Larval desiccation tolerance in the parental lines (A), and in their F_1_ hybrids with the *ru cu ca* marker strain (B)

To test whether plasticity of recombination rate and crossover interference shows fitness dependence, we compared the following hypotheses: (a) H_0_ - no difference in the considered recombination parameters between the treated and non-treated F1 females in both the *S*- and *T*-lines; (b) H_1_ – no difference between the treated and non-treated *T*-lines; (c) H_2_ – no difference between the treated and non-treated *S*; and (d) no restriction on parameters of the two groups of lines, i.e. assuming that both lines may display treatment-induced changes.

With respect to the plasticity of *recombination rate*, we found that for two out of five examined intervals (*h-th* and *cu-sr*), H_1_ but not H_2_ significantly differed from H_0_ (*P*=1.45×10^−3^ and 6.51×10^−3^, respectively; after correction for multiple tests, *P*=0.024 and 0.039, respectively). A slight and non-significant desiccation-induced increase in recombination rate was also observed in the *T*-lines across both intervals. When considered jointly, induced changes in both groups of the lines give rise to a two-fold improvement of statistical significance for the recombinogenic effect of desiccation (H_3_ vs H_0_). For three other intervals no effect of treatment on recombination rate was observed (i.e., none of the hypotheses, H_1_, H_2_, or H_3_, significantly differed from H_0_) (Table 1).

**Table 1.**
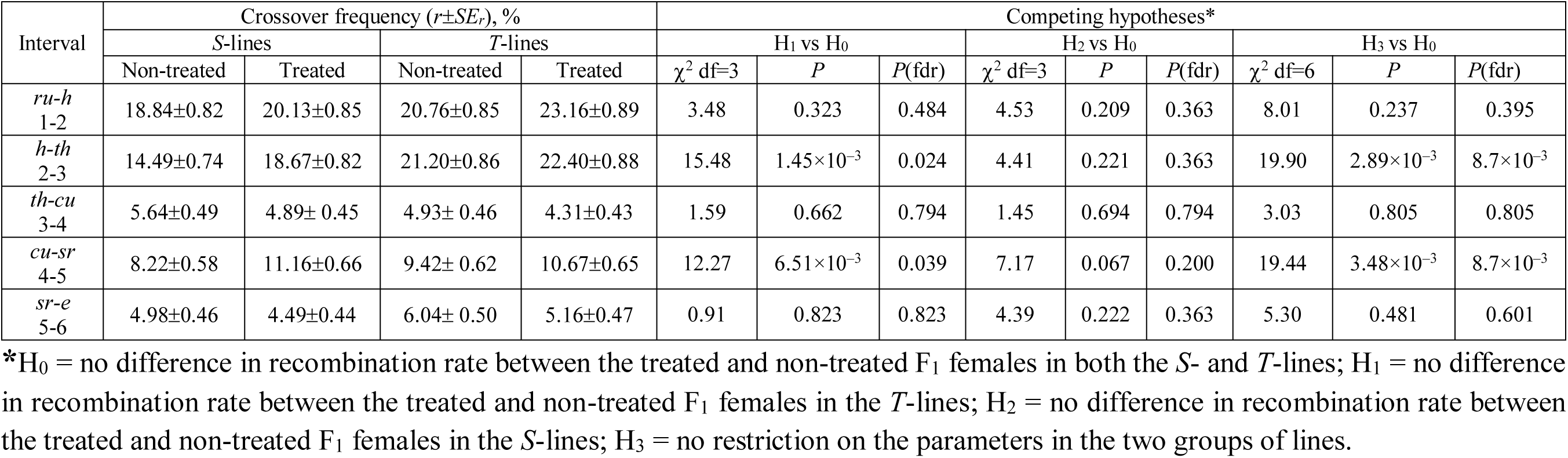
The effects of desiccation on recombination rates in chromosome 3 in desiccation-sensitive and desiccation-tolerant *Drosophil* lines

| Interval | Crossover frequency ($r \pm SE_r$), % | | | | Competing hypotheses* | | | | | | | | |
| --- | --- | --- | --- | --- | --- | --- | --- | --- | --- | --- | --- | --- | --- |
| | *S*-lines | | *T*-lines | | $H_1$ vs $H_0$ | | | $H_2$ vs $H_0$ | | | $H_3$ vs $H_0$ | | |
| | Non-treated | Treated | Non-treated | Treated | $\chi^2$ df=3 | *P* | *P*(fdr) | $\chi^2$ df=3 | *P* | *P*(fdr) | $\chi^2$ df=6 | *P* | *P*(fdr) |
| *ru-h*<br>1-2 | 18.84±0.82 | 20.13±0.85 | 20.76±0.85 | 23.16±0.89 | 3.48 | 0.323 | 0.484 | 4.53 | 0.209 | 0.363 | 8.01 | 0.237 | 0.395 |
| *h-th*<br>2-3 | 14.49±0.74 | 18.67±0.82 | 21.20±0.86 | 22.40±0.88 | 15.48 | $1.45 \times 10^{-3}$ | 0.024 | 4.41 | 0.221 | 0.363 | 19.90 | $2.89 \times 10^{-3}$ | $8.7 \times 10^{-3}$ |
| *th-cu*<br>3-4 | 5.64±0.49 | 4.89±0.45 | 4.93±0.46 | 4.31±0.43 | 1.59 | 0.662 | 0.794 | 1.45 | 0.694 | 0.794 | 3.03 | 0.805 | 0.805 |
| *cu-sr*<br>4-5 | 8.22±0.58 | 11.16±0.66 | 9.42±0.62 | 10.67±0.65 | 12.27 | $6.51 \times 10^{-3}$ | 0.039 | 7.17 | 0.067 | 0.200 | 19.44 | $3.48 \times 10^{-3}$ | $8.7 \times 10^{-3}$ |
| *sr-e*<br>5-6 | 4.98±0.46 | 4.49±0.44 | 6.04±0.50 | 5.16±0.47 | 0.91 | 0.823 | 0.823 | 4.39 | 0.222 | 0.363 | 5.30 | 0.481 | 0.601 |

\* $H_0$ = no difference in recombination rate between the treated and non-treated $F_1$ females in both the *S*- and *T*-lines; $H_1$ = no difference in recombination rate between the treated and non-treated $F_1$ females in the *T*-lines; $H_2$ = no difference in recombination rate between the treated and non-treated $F_1$ females in the *S*-lines; $H_3$ = no restriction on the parameters in the two groups of lines.

With respect to the plasticity of *crossover interference*, the effect held for the 3L but not for the 3R arm: for two out of the four examined pairs of adjacent intervals (*ru-h-th* and *h-th-cu*), H_1_ but not H_2_ significantly differed from H_0_ (*P*=1.43×10^−2^ and 2.72×10^−3^, respectively; after correction for multiple tests, *P*=0.064 and 0.024, respectively). For two other pairs, neither H_1_ nor H_2_ significantly differed from H_0_ (Table 2). Given that two intervals (*th–cu* and *sr*–*e*) are small, rates of double crossovers with their adjacent intervals are expected to be low. Thus, we additionally examined desiccation-induced changes in interference in some ‘derivative’ intervals (Table 3). Overall, the difference in interference response to treatment between the *S-* versus *T*-lines was relatively robust. Similar to the interval pair *h-th-cu*, the derivative pair *h-th-sr* that includes the same left-arm interval *h-th*, but additionally extends onto the centromeric interval *th-sr*, desiccation treatment caused an increase in the double-crossover rate (relaxation of positive interference) in the *S*-, but not in the *T*-lines. Moreover, we observed a significant *reduction* in the double-crossover rate in the *T*-lines. Taken together, these opposite-direction changes are manifested in highly significant rejection of H_0_ in favor of H_3_. The next two pairs of derivative intervals, *h-cu-sr* and *h-cu-e*, with the centromere in the left interval, demonstrates a similar pattern: a treatment-induced increase in double-crossover frequency in the *S*-lines and simultaneous decrease in the *T*-lines. Unlike the above-considered effects of treatment on crossover interference in the pericentromeric/left-arm pairs of intervals, no significant changes were observed in analogous combinations of pericentromeric/right-arm pairs of intervals.

**Table 2.**
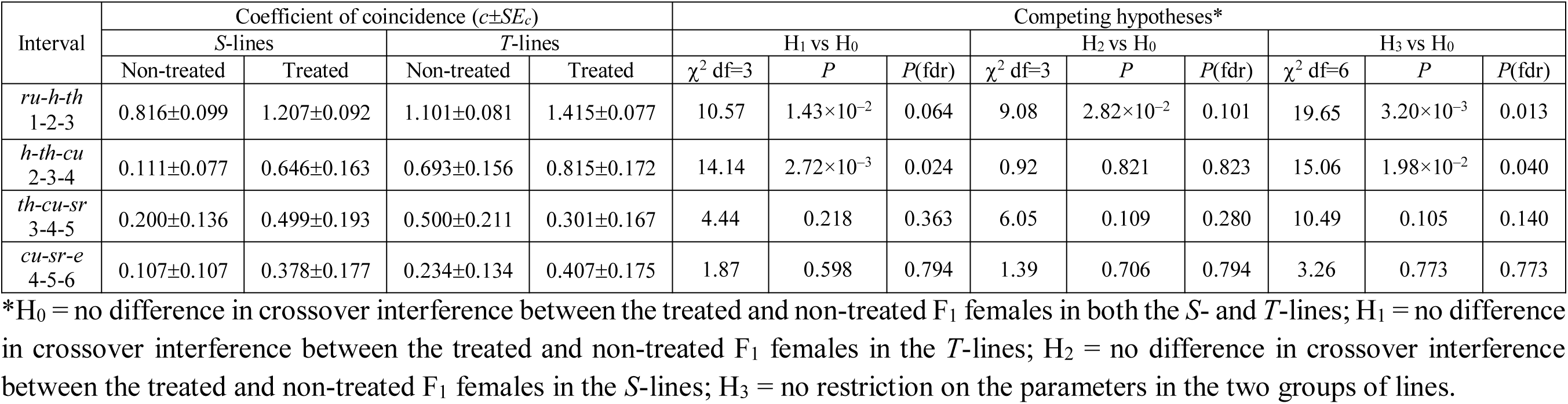
The effect of desiccation on crossover interference in chromosome 3 in desiccation-sensitive and desiccation-tolerant *Drosophila* lines

**Table 3.**
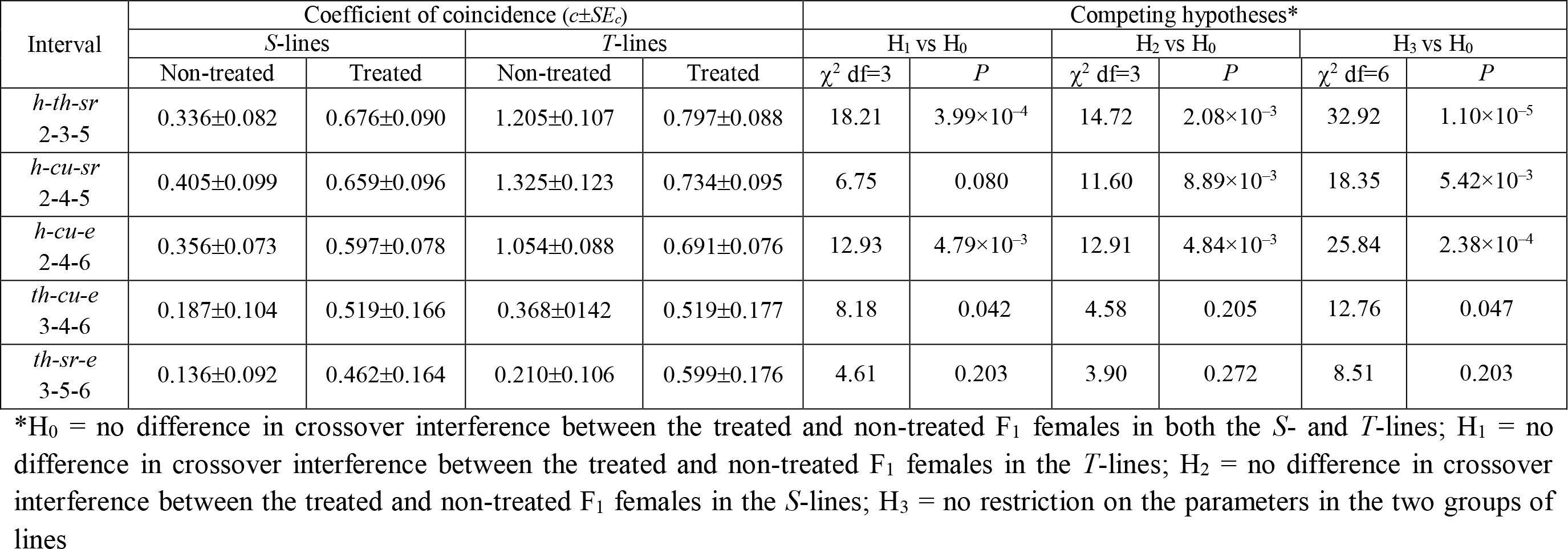
The effect of desiccation on crossover interference in chromosome 3 in desiccation-sensitive and desiccation-tolerant *Drosophila* lines: derivative intervals

In general, our results indicate that desiccation stress affects both recombination rate and crossover interference. At that, wherever recombination rate and interference are plastic, higher reactivity is associated with lower fitness. Remarkably, in certain chromosomal regions, stress-tolerant genotypes demonstrate a tendency to decrease the double-crossover rate upon treatment, exactly opposite to that shown by stress-sensitive genotypes.

Given the segment-specific recombination response to desiccation treatment, a question arises if the reacting intervals carry genes involved in desiccation tolerance. Indeed, we find the two intervals to be enriched in functional terms related to stress response, including chitin metabolic processes, oxidative stress, and transmembrane transport, among others (Table 4). We also examined the intervals for any major genome sequence changes that may have been driven by long-term (48 generations) selection for desiccation tolerance (previously described by Aggarwal et al. 2015). Our earlier comparative analysis of whole genome sequence data on the *S*- and *T*-lines (Kang et al. 2016) revealed only one hard-sweep candidate region in 3L (11612902-11960953), residing in the *h–th* interval. In addition, out of the 17 potential soft-sweep regions detected in 3L, five were within the *h–th* interval, and two more were found at the opposite ends of the interval. In 3R, the only hard-sweep region (15032772–15970755) was found in the *cu–sr* interval, and out of 14 potential soft-sweep regions, two (adjacent) regions were located within the *cu–sr* interval. A total of 64 and 44 non-synonymous substitutions in the coding regions mapped to the *h–th* and *cu–sr* intervals, respectively. Out of these, 26 genes in the *h–th* interval were concentrated in three large “islands” ((i) from the *klu* gene to *CG34012*; (ii) from *Hip1* to *CG34428*, and (iii) from *Hsc70Cb* to *RecQ5*). The *SuUR* gene from island (i), *CG10948* and *CG11267* from island (ii), and *fz* from island (iii) were earlier reported as down-regulated in a study on responses to desiccation in natural populations of a desert drosophilid (Rajpurohit et al. 2013). In the *cu-sr* interval, only one sweep region was found (between *Irc* and *Mur89F* genes). Inside this region, *Mur89F* was earlier reported to be up-regulated and *cal1* was down-regulated in response to desiccation (Rajpurohit et al. 2013). Although the resolution (marker density) of the current study is insufficient to provide more insights into potential associations between the interval-specific recombination responses and the functions of genes residing in the intervals, these tentative comparisons are promising and initially suggestive of such associations.

**Table 4.**
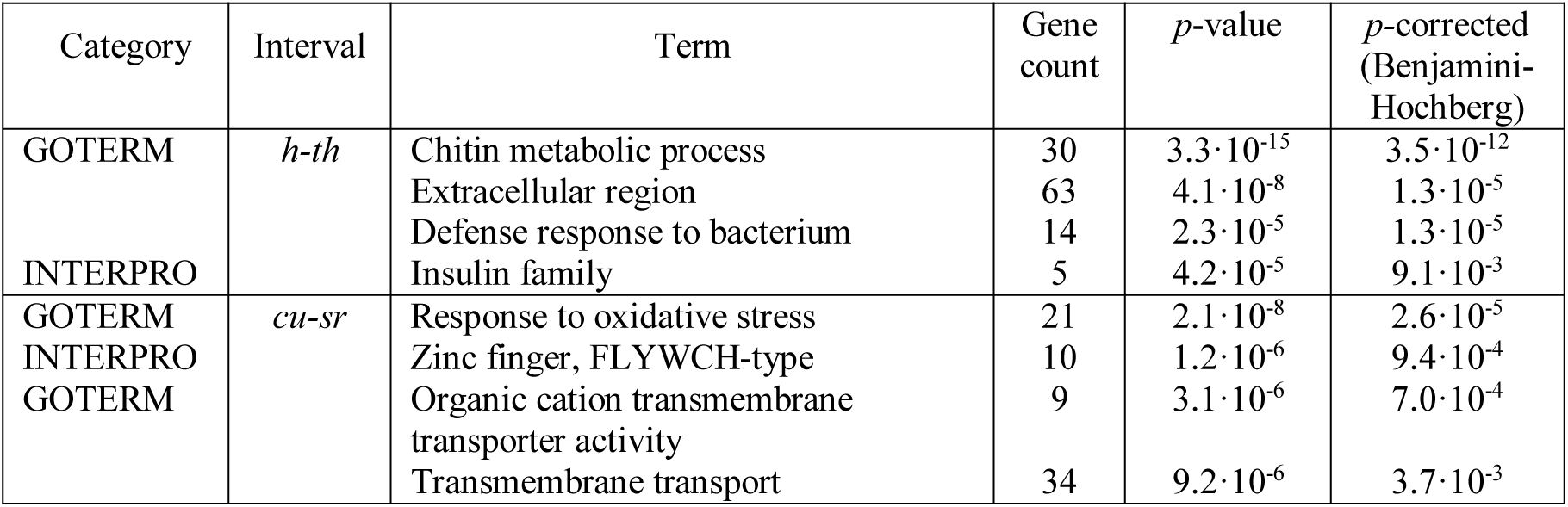
Gene Ontology enrichment tests for genes found in the *h-th* and *cu-sr* genomic intervals: the responding intervals *of* chromosome 3 were contrasted with all *D. melanogaster* genes using *DAVID* (shown are only terms with *p*-value <0.01 after the Benjamini-Hochberg correction)

## DISCUSSION

### Desiccation as a recombinogenic factor

In the *S*-lines, desiccation significantly raised crossover frequency in two out of the five examined intervals: *h-th* and *cu-sr*. Both intervals are close to the centromere; recombination in such regions is long known to be highly regulated, as well as highly reactive with respect to environmental stressors (Plough 1921; Grell 1978). Our results indicate that desiccation is a recombinogenic factor; to our knowledge, this has been shown earlier only in maize (Verde 2003).

In the *S*-lines, desiccation caused a significant increase in double-crossover rates in two of the four examined pairs of intervals: *ru-h-th* and *h-th-cu*. This result is consistent with a heat-induced relaxation of interference observed earlier in *Drosophila* chromosomes X, (Hayman and Parsons 1960; Grell 1978), 2 and 3 (Grell 1978). Whether or not changes in interference are necessarily coupled with those in recombination has been a hot topic discussed for decades (Grell 1978; Denell and Keppy 1979; Foss et al. 1993; Fujitani et al. 2002; Zhang et al. 2014; Aggarwal et al. 2015; Zickler and Kleckner 2016). Indeed, higher recombination rates are often, but not always, associated with relaxation of positive interference and even appearance of negative interference (recently reviewed in Aggarwal *et al.*, 2015). Herein observed increase in double-crossover rates in the *S*-lines did not strictly coincide with an increase in crossover frequencies in one or both adjacent intervals, consistent with results from some earlier studies (Grell 1978; Denell and Keppy 1979). Thus, we do not automatically derive the former from the latter. Yet, we consider changes in recombination and interference to be of the same nature, namely as manifestations of meiosis deregulation. Similarly, meiotic mutants with deregulated recombination may show a more uniform distribution of crossover frequency along chromosomes and a relaxation of positive crossover interference and crossover homeostasis compared to the wild type genotypes (Baker and Hall 1976; Szauter 1984; Zhuchenko and Korol 1985; Zetka and Rose 1995; Séguéla-Arnaud et al. 2015). However, this conclusion is based mainly on measurements of recombination/interference parameters and mechanistic considerations rather than on direct tests of changes in their variation.

### Fitness dependence of desiccation-induced changes in recombination rate and interference

The herein revealed difference between the *S*- and *T*-lines in the plasticity of recombination rates under desiccation is consistent with results of other studies, in which fruit flies were exposed to other environmental stressors. Thus, Kilias *et al.* (1979) observed higher crossover frequencies in a *Drosophila* line with low activity of alcohol dehydrogenase compared to another, high-activity line. The authors suggested that the former strain had lower fitness due to higher sensitivity to alcohol intoxication. However, given that alcohols are capable to cause DNA damages, it is impossible to distinguish between the stressor effect on somatic fitness and its direct effect on germline DNA metabolism. Zhuchenko and Korol (1985) assessed the effect of heat treatment on recombination rate in the pericentromeric interval *b-cn* of chromosome 2 in heterozygotes of crosses between a marker line and 12 *Drosophila* lines varying in thermotolerance. They found a three-fold increase in the variance of *r*_*b-cn*_ due to the treatment and a significant negative correlation between the heat-induced changes in recombination rate and the corresponding changes in fertility and fecundity. Then Korol *et al*. (1994) revealed that heterozygous heat-sensitive *Drosophila* males showed, when heat-shocked, a several-fold higher increase in crossover frequency in chromosome 2 compared to heat-tolerant ones. Yet, recombination reported therein could be both meiotic and pre-meiotic (Hiraizumi 1971; Woodruff and Thompson 1977). Finally, fitness dependence of stress-induced changes in meiotic recombination was demonstrated by Zhong and Priest (2011). In their experiments, heat-treated *Drosophila* females showed an increase in crossover frequency that negatively correlated with their heat-tolerance measured as productivity.

Another negative association between recombination rates and individual fitness was recently reported by Jackson *et al*. (2015). However, their conclusion was based on examining a single genotype, as so that variation in recombination rate was observed within the reaction norm. Hunter *et al*. (2016) analyzed a vast set of genotypes (~200 lines from the Drosophila Genetic Reference Panel) and reported a negative correlation between crossover rate in the *y–v* interval (X-chromosome) and flies’ response to certain chemicals. One of them, paraquat, is a model inducer of oxidative stress; thus, paraquat tolerance indeed seems an adequate measure for fitness. However, the phenotype traits were measured in the lines themselves, while recombination rates – in their F_1_ hybrids with marker stocks; thus, the obtained correlation values may have been biased.

We interpret the difference in recombination response to desiccation between our *S*- and *T*-lines as evidence for fitness-dependent recombination plasticity. An alternative explanation might be that recombination rates, observed in the *T*-lines, might have already approached their limit and cannot be increased much further. However, in the *S*-lines, desiccation stress raised recombination rates in some intervals to a level even higher than the average of the *T*-lines (e.g. in case of *cu*–*sr* interval). Dissecting the association between the plasticity of recombination rate and of crossover interference on one hand, and fitness on the other, will require further tests with a considerably higher number of intervals and examined genotypes.

Physiological and molecular mechanisms responsible for fitness-dependent plasticity of recombination remain underexplored. We did not investigate these processes but the current study will serve as a necessary stepping stone for future mechanistic approaches; nevertheless, some speculations regarding possible mechanisms are presented below. Theoretically, higher stress tolerance of soma and of germline may develop in a more or less parallel way. If so, the observed difference between the *S*- and *T*-lines in terms of their recombination response to stress merely reflects the intrinsic capabilities of the germline cells. Alternatively (which is our hypothesis), the difference originates, at least partially, as a transition of stress effects from somatic cells to the germline. Evidence for such soma-to-germline signaling has indeed begun to emerge. For example, somatic status has been known to affect crucial stages of germline development in *Drosophila*, including sex determination, gonad differentiation, and apoptosis. Regulatory signals may originate from adjacent or even distant tissues exposed to various stress effects, e.g., such as malnutrition (Laws and Drummond-Barbosa 2017). Thus, strong germline effects have also been found in fruit flies subject to behavioral stress, such as predator presence and even communication with individuals previously exposed to predators (Kacsoh et al. 2015). These observations indicate that integrated, systemic soma-to-germline signaling may indeed exist. Such signaling may be based, for example, on the interaction between ecdysone/let-7 pathway in soma and Wnt pathways in germline (Fagegaltier et al. 2014; König and Shcherbata 2015). Interestingly, growing evidence indicates that soma-germline communication is likely to be reciprocal (Parisi et al. 2010). All the above suggests that soma and germline tightly communicate, which allows transmitting signals (including stress-associated ones). We believe that this communication might contribute to fitness-dependent regulation of recombination.

Unraveling the potential role of behavioral (neurogenic) stresses in the regulation of germline processes, including meiotic recombination, in species with the highly organized nervous system, like mammals, is of particular interest. In a pioneering research program in the 1980s, Borodin and Belyaev provided unique empirical evidence that emotional stress increases meiotic recombination rate in house mouse. Specifically, mouse males exposed to severe overcrowding displayed significantly higher recombination rates in chromosome 1 and 2 (Borodin and Belyaev 1980a, b). A considerable increase in the rates of XY and autosomal univalents in metaphase 1 and aneuploidy in metaphase 2 under acute immobilization stress was also recorded (Gorlov and Borodin 1986). Furthermore, a reduction in DNA repair synthesis was detected in the spermatids of the treated males (Borodin 1987). The availability of new genomic techniques provides an exciting opportunity to explore these patterns in conjunction with genome information.

### Evolvability of fitness dependence

If fitness-dependent recombination confers some benefits, it may evolve as an adaptive trait. To model the evolution of fitness-dependent recombination, one can utilize the standard model for the evolution of selectively neutral locus modifying recombination rates (Kimura 1956; Nei 1967). Within this modifier framework, two forces are discussed in the literature as being capable (at least potentially) of driving the evolution of fitness-dependent recombination. The first force is related to benefits that accrue from a recombination-modifying allele capable to affect its own linkage to the selected locus. Specifically, a modifier allele that through recombination tends to abandon the linkage with the unfavorable allele of the selected locus will spread owing to the “right” association. This mechanism is now commonly referred to as the “abandon-ship” model (Agrawal et al. 2005). The second force is related to the benefits from the plasticity of recombination rates *between the selected loci*. Such benefits can arise from protecting good selected haplotypes (i.e., by decreasing recombination rate under high fitness), while producing novelty by utilizing the poor ones (i.e., increasing recombination under low fitness). A modifier allele underlying this strategy can spread using associations with favorable combinations of the selected alleles.

In haploids, each of these two forces alone can drive the evolution of fitness-dependent recombination (Gessler and Xu 2000; Hadany and Beker 2003a, b; Agrawal et al. 2005; Wexler and Rokhlenko 2007). In diploids, the “abandon-ship” mechanism has been shown to be inefficient (Agrawal et al. 2005). However, our recent simulations demonstrate that fitness-dependent recombination can evolve in diploids under certain scenarios, such as cyclical selection (Rybnikov et al. 2017), Red Queen dynamics (Rybnikov et al. 2018a), and mutation-selection balance (Rybnikov et al. 2018b). In all our models, fitness-dependence was assumed only for recombination within the selected system. This can make plastic recombination beneficial only if there is a variation in fitness among heterozygous genotypes, which requires at least three selected loci. Importantly, the evolutionary advantage/disadvantage of fitness-dependent recombination is determined by a trade-off between two opposite effects: benefits from protecting best allele combinations, and costs of shifting population mean recombination rate (Rybnikov et al. 2018b). The cost often outbalances the benefits, which may be one of the reasons why some experimental-evolution studies reported no selection for recombination rate plasticity (Kerstes et al. 2012; Kohl and Singh 2018).

Fitness-dependent interference may be subject to indirect selection as well. Its evolvability has been demonstrated in a modifier model by Goldstein *et al*. (1993). Recent theoretical models also support the evolvability of fitness dependence of other processes affecting genetic variation: sex (Hadany and Otto 2007), mutation (Shaw and Baer 2011), and dispersal (Gueijman et al. 2013). This tempts us to conclude that the evolvability of fitness dependence may be a widespread phenomenon in nature. The consistent evolutionary pattern is that generating *de novo* or releasing standing genetic variation is modulated by individual fitness so that less fitted individuals tend to produce more variable progeny (Korol et al. 1994). Ervin Bauer put forward in a somewhat paradoxical form a similar idea that ‘losers’ rather than ‘winners’ in the struggle for existence provide the raw material for evolutionary change (cited according to (Korol et al. 1994)). The general reason for such an increased production of genetic variability can also be seen in the ‘genomic stress’ caused by external and internal factors (McClintock 1984; Hoffmann and Parsons 1991; Korol et al. 1994; Thaler 1994). How common is such a system of variability control, with a reverse modulating effect of fitness, remains an open question deserving further studies.

## ACKNOWLEDGMENTS

We are grateful to two anonymous reviewers for critical comments and corrections which allowed to improve the manuscript. We also thank Kostas Illiadi, Eugene Kandel and Marios Kyriazis for fruitful discussions on soma-germline interactions.

## FUNDING

The study was supported by the Israel Science Foundation (grant 1844/17); the University Grants Commission, India (DS Kothari project grant F.4-2/2006(BSR)/BL/16-17/0330); the Council for Higher Education of the Israeli Ministry of Education; and the Israeli Ministry of Aliyah and Integration.

